# From Blue to Pink: Resazurin as a High-Throughput Proxy for Metabolic Rate in Oysters

**DOI:** 10.1101/2025.11.06.686367

**Authors:** Ariana S. Huffmyer, Noah Ozguner, Madeline Baird, Colby Elvrum, Carolyn Kounellas, Dash Dicksion, Samuel J. White, Louis Plough, Mackenzie R. Gavery, Noah Krebs, William Walton, Jessica Small, Madeline Pitsenbarger, Healy Ealy-Whitfield, Steven Roberts

**Affiliations:** University of Washington, School of Aquatic and Fishery Sciences, 1122 NE Boat St Box 355020 Seattle, WA 98195; United States Department of Agriculture, Agricultural Research Service, Pacific Shellfish Research Unit, 2030 SE Marine Science Drive, Newport, OR 97365; Environmental and Fisheries Sciences Division, Northwest Fisheries Science Center, National Marine Fisheries Service, National Oceanic and Atmospheric Administration, 2725 Montlake Blvd E, Seattle, WA 98112; Virginia Institute of Marine Science, William & Mary, 1370 Greate Road, Gloucester Point, VA 23062

**Keywords:** *Crassostrea gigas*, *Crassostrea virginica*, metabolism, thermal stress, shellfish, resilience

## Abstract

Metabolic rate assays are critical tools for assessing organismal stress and resilience, yet their widespread application in aquaculture and ecological monitoring is limited. Improving these assays is essential for hatchery managers, farmers, and scientists seeking to identify resilient stocks and monitor stress in shellfish populations. Resazurin, a redox-sensitive dye commonly used in cell viability assays, offers a promising, high-throughput assay for metabolic rate assessment, but its application at the whole-organism level remains under explored. This study evaluates the efficacy of a resazurin-based metabolic assay in oysters (*Crassostrea gigas* and *Crassostrea virginica*) through four experimental approaches: (1) adaptation of the resazurin assay to measure oyster metabolism, (2) examination of temperature effects on oyster metabolism, (3) characterization of acute thermal stress responses, (4) examination of genetic variability in metabolism, and (5) correlations between metabolism and predicted performance in a selective breeding case study. Our findings confirm that resazurin fluorescence is correlated with oxygen consumption, validating its use as a measure of metabolism. Thermal performance assays reveal expected metabolic responses to temperature, including identification of optima and tipping points where metabolic stimulation shifts to depression under temperature stress. Acute thermal stress experiments demonstrate that oysters exhibiting greater metabolic depression are more likely to survive, supporting metabolism as a predictor of mortality. Further, genetic variation in stress responses is detected as family-level variation in metabolism. Metabolism of 50 families (*C. virginica*) selectively bred for performance in varying environments was measured and significantly correlated with predicted performance. By establishing resazurin as an additional reliable and scalable method for metabolic assessment, this study lays the groundwork for its broader adoption in aquaculture and conservation. Implementing this approach may provide a tool to enhance stock selection, improve hatchery management practices, and support adaptive strategies in the face of climate variability and increased environmental stress in coastal oceans.

## Introduction

Metabolic assays are powerful tools for assessing physiological stress and resilience in marine organisms, particularly as climate change and environmental stressors increasingly threaten aquatic ecosystems. By quantifying energy expenditure, these assays provide critical insight into organismal health, performance, and survival potential (Sokolova, 2013). In shellfish, metabolic measurements are essential for understanding responses to environmental fluctuations such as temperature extremes, hypoxia, and ocean acidification (Chen et al., 2022; Lannig et al., 2010, 2006; Le Moullac et al., 2007; Méthé et al., 2020; Widdows et al., 1989). There is a need to develop rapid, high-throughput metabolic assays for use by researchers and aquaculture practitioners to enhance efforts to monitor stock health, identify resilient genotypes, and optimize hatchery management.

Resazurin, a redox-sensitive dye, presents a promising assay for high-throughput metabolic rate assessments. Widely used in cell viability and toxicity assays due to its simplicity, sensitivity, and cost-effectiveness (Petiti et al., 2024), resazurin provides a fluorescent indicator of metabolic activity. As the organism conducts metabolism, resazurin undergoes a stepwise reduction from its blue, non-fluorescent, oxidized form to pink, fluorescent, resorufin by NADH and reductases, producing a strong fluorescent response (Candeias et al., 1998; Chen et al., 2018; O’Brien et al., 2000). This fluorometric shift provides a robust, quantitative measure of whole-organism metabolic activity. Resazurin-based assays have been extensively applied across diverse fields, including cytotoxicity screening (Fields and Lancaster, 1993; Hussain et al., 2011; O’Brien et al., 2000; Pace and Burg, 2015; Petiti et al., 2024), microbial metabolism studies (Fai and Grant, 2009; González-Pinzón et al., 2012; Van den Driessche et al., 2014; Zare et al., 2015), and biomedical applications (Anoopkumar-Dukie et al., 2005; McMillian et al., 2002). In marine biology, resazurin has been used to assess sperm viability in marine invertebrates and fish (Hamoutene et al., 2000) and to measure metabolic activity in zebrafish larvae (Reid et al., 2018; Renquist et al., 2013), demonstrating its potential as a non-lethal, reliable metabolic indicator. One study applied resazurin to evaluate hemocyte viability in oysters exposed to toxins (Estrada et al., 2021), but its use for whole-organism metabolic rate assessment in shellfish remains largely unexplored.

Despite its promise, adapting and testing resazurin for metabolic rate assays has not yet occurred in shellfish and requires testing of factors such as incubation time, stress exposure profiles, and comparison with established methods. By addressing these technical considerations, resazurin-based assays could offer a scalable, complementary approach for evaluating metabolic responses in shellfish, with applications in both research and aquaculture to predict and evaluate stress resilience and performance. Therefore, the objective of this study was to develop and validate a resazurin-based metabolic assay for oysters (*Crassostrea gigas* and *Crassostrea virginica*), focusing on its utility for assessing metabolic resilience to environmental stress.

We had five objectives in this study: (1) Adaptation of the resazurin assay to measure whole organism oyster metabolism. In this objective, we expected resazurin fluorescence measurements of metabolism to positively correlate with oxygen consumption. Further, we expected resazurin fluorescence to increase with oyster size and to measure metabolism in live oysters higher than that of empty shell and blank samples. (2) Examination of temperature effects on oyster metabolism. In this approach, we hypothesized that temperature would affect metabolism with metabolism peaking at thermal optima and decreasing at high temperatures. (3) Characterization of acute thermal stress responses. We expected to identify clear signals of acute stress in metabolism with metabolism corresponding to mortality. (4) Examination of genetic variability in metabolic responses. We hypothesized that families would exhibit distinct metabolic responses to thermal stress. (5) Conduct a case study to identify relationships between metabolism and performance. We expected that metabolism would correlate with predicted performance breeding values in selectively bred *C. virginica* families.

By establishing a high-throughput metabolic assay for shellfish that is informative for stress resilience and performance assessments, this study provides a new application of a practical tool for monitoring shellfish health, identifying stress-resilient lineages, and improving aquaculture management in a changing climate.

## Materials and Methods

We conducted a series of experiments in this study to adapt the resazurin assay for use in oysters and test protocols to evaluate oyster stress response and performance. First, we describe the general protocol for resazurin metabolic assays and then provide methodologies for each objective individually.

### A. Resazurin standard protocol

We have included a general resazurin assay protocol for use in shellfish in **Supplementary Information Appendix A.** All experiments used a standardized resazurin solution. The resazurin assay protocol is based on previously published protocols in zebrafish (Reid et al., 2018; Renquist et al., 2013). A concentrated resazurin stock solution was prepared by dissolving 1.0 g of resazurin sodium salt (Cat. R7017 Sigma-Aldrich, Saint Louis, MO) in 20 mL (4.76% w/v) of deionized (DI) water with 20 µL dimethyl sulfoxide (DMSO; 0.10% v/v). The concentrated stock solution was used to generate a working solution for assays and stored in the dark at 4°C until use. Fresh working solution (111 µg/mL resazurin) was prepared prior to each experiment by mixing 0.22% (v/v) resazurin stock solution, 0.1% DMSO (v/v), and 1.0% (v/v) antibiotic antimycotic 100x Penn/Strep/Fungizone solution (Cat. SV30079.01, Cytiva, Marlborough, MA) in filtered seawater or DI water adjusted to 25 ppt using Instant Ocean salts (Instant Ocean, Blacksburg, VA). For example, a 500 mL working solution included 494 mL seawater, 1.11 mL resazurin stock solution, 500 µL DMSO, and 5 mL antibiotic solution. The working solution was either used immediately or stored at 4 °C in the dark and used within 1 week.

Oysters were then photographed to measure shell length (mm) in ImageJ (Rueden et al., 2017) for size normalization and then allocated to individual size appropriate containers for incubations. For example, small seed (3-8 mm) were added into 96-well plates with larger seed (6-12 mm) added to 48 or 24-well plates. Larger oysters (8-20 mm) can be added to 12 or 6-well plates with increasing sizes (>20 mm) added to individual plastic cups (20-40 mL). Containers were then filled with the prepared resazurin solution with empty containers serving as resazurin blank samples.

Samples were read on a plate reader in fluorescence mode with emission of 530 nm and excitation of 590 nm set to collect endpoint measurements reading from the top of the sample. For samples in multi-well plates, plates were placed directly on the plate reader for readings while oysters in larger cups were measured by taking a 200 µL subsample and adding to a 96-well plate for readings. Following the initial measurement, oysters were exposed to designated temperature conditions controlled by benchtop incubators with readings collected every subsequent hour for 4-6 h. Specific equipment and sample frequency are described for each objective below. See an example of resazurin measurements in 96-well plates in **Fig 1**.

**Fig 1.**
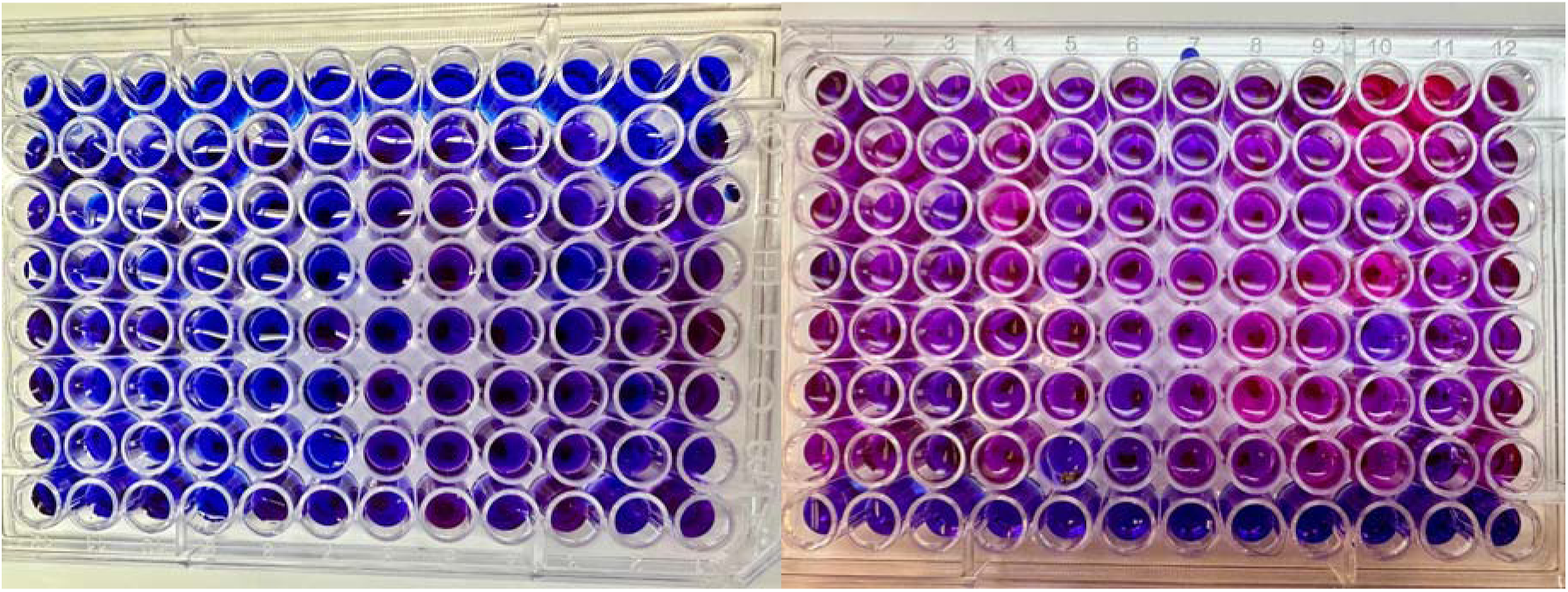
Visual examples of resazurin assays conducted on oysters. Note that brighter pink colors indicate higher metabolism as resazurin is reduced to resorufin. The left photograph shows an example of oysters in 96-well plates early in an incubation (low fluorescence) and the right photograph shows an example of oysters in 96-well plates late in an incubation (high fluorescence). Note the bottom most row includes blank wells.

Fluorescence measurements were calculated as fold change by normalizing to the initial fluorescence value and then corrected for background changes in resazurin solution fluorescence by subtracting mean fluorescence values of blanks for each replicate plate or group. Corrected fluorescence values were then normalized to size by dividing by shell length to generate size-normalized fold change in fluorescence. Hereafter, size-normalized fold change in fluorescence is referred to as “metabolism”. Metabolism was quantified as metabolism over time and evaluated by calculating “total metabolic activity” as Area Under the Curve (AUC) using trapezoidal integration in the *pracma* package (Borchers, 2023). The effect of experimental variables on metabolism and total metabolic activity was then tested using analysis of variance (ANOVA) or linear mixed effect models in the *lme4* package (Bates et al., 2014) with post hoc comparisons using estimated marginal means tests in the *emmeans* package (Lenth, 2018). All analyses were conducted in R Statistical Programming v4.4.1 (R Core Team, 2022). Specific statistical methods are described below for each objective.

### B. Objective 1: Adaptation of resazurin assay to measure whole organism oyster metabolism

#### (1) Correlations between resazurin and oxygen measurements

We conducted oxygen consumption and resazurin assays on seed to examine the relationship between the two methods for measuring metabolism on the same individual oyster following the standard resazurin protocol described above. We selected n=95 oyster seed (4-7 mm) obtained from Pacific Hybreed (Port Orchard, WA) and held in tanks supplied with water adjusted to 25 psu using Instant Ocean salts (Instant Ocean, Blacksburg, VA) at the University of Washington. Oxygen measurements were conducted first on all individuals using a 24-well Loligo (Viborg, Denmark) glass plate (1.7 mL well volume) with PreSens (Regensburg, Germany) sensor spots (PSt5-2334-01) in the MicroResp software (v1.2.1.0). Microplate respirometry measurements have been reported in shellfish previously (Gurr et al., 2025, 2024). Wells were loaded with air saturated 0.2 µm filtered seawater (FSW, 25 psu) and sealed with plate sealing strips. Each run contained n=20 oysters per plate with n=4 blank wells containing only FSW. Measurements were conducted at 27°C with oxygen concentration (mmol per L) measured every 15 s for 50 min to 1 h. Five plates were run on the same day to measure all individuals. Oxygen consumption rates were extracted from oxygen measurements using localized linear regressions in the *LoLinR* package (Olito et al., 2017) using the percentile rank method with alpha set to 0.4 in R. Oxygen consumption rates were corrected by subtracting blank well values and dividing by shell length to produce µmol O_2_ mm^-1^ min^-1^.

Resazurin measurements were then conducted on the same set of individuals the following day in 48-well plates (n=3; 1 mL well volume) with n=40 oysters and n=8 blanks per plate. Measurements were conducted at 27°C over a 5 h period with hourly measurements using a BioTek Synergy HTX multi-mode plate reader (Agilent, Santa Clara, CA) in fluorescence mode with emission at 530 nm (528/20 nm filter) and excitation at 590 nm (590/20 nm filter) with associated software (Agilent Gen5 Software). Metabolism (fold change in fluorescence mm^-1^) during the first hour of incubation was used to directly compare metabolism via resazurin with oxygen consumption. We then compared oxygen (cube-root transformed) and resazurin measurements (log-transformed) using a Pearson correlation. There was no observed mortality.

#### (2) Fluorescence signals from shells compared to live animals

In order to confirm resazurin signals are due to live animal metabolism and not an artifact of microbial communities on shells, we examined the difference in metabolism of live oysters (15-20 mm; n=15), empty shells (15-20 mm; n=15), and blank wells (n=6) of resazurin solution using 12-well plates at 35°C hourly for 4 h using a BioTek Synergy HTX multi-mode plate reader (Agilent, Santa Clara, CA). There was no observed mortality during the trials. We then examined metabolism before and after size normalization to compare fluorescence between blanks, empty shells, and live oysters. Differences were tested using one-way ANOVA tests with sample type as the main effect.

#### (3) Relationship between oyster size and metabolism

We evaluated the relationship between total metabolism, calculated as AUC (prior to size normalization), and oyster size by combining metabolism measurements and shell length (mm) data obtained from objectives 3 and 4 (described below) because these experiments were conducted at the same site (University of Washington) with a similar size class (approx. 15-35 mm shell length) at 18°C. The relationship between oyster size and metabolism was evaluated using a Spearman correlation test as well as a linear model (LM; square-root transformed AUC).

### C. Objective 2: Examination of temperature effects on oyster metabolism

#### (1) Thermal response in small seed (3-8 mm)

We examined the effect of temperature on metabolism measured using the resazurin fluorescence assay by exposing small seed to a range of temperature conditions in two tests of small seed (3-8 mm) and larger seed (5-15 mm). First, we exposed small seed (3-8 mm shell length) to 20, 36, 38, 40, and 42°C for 4 h. Seed were added to individual wells of 96-well plates filled with 280 µL of resazurin working solution. Oyster seed were obtained from the Point Whitney Jamestown S’Klallam Shellfish Hatchery from two separate broodstock spawning groups and held at 14°C in two tanks filled with water adjusted to 25 psu using Instant Ocean salts (Instant Ocean, Blacksburg, VA) at the University of Washington. Prior to measurements, seed were photographed to measure shell length (mm) in ImageJ (Rueden et al., 2017). Fluorescence was measured by placing the plate containing oysters directly on a plate reader (BioTek FLx800; Agilent, Santa Clara, CA) in fluorescence mode with emission of 530 nm (530/20 nm filter) and excitation of 590 nm (590/20 nm filter) with associated software (BioTek Gen5). Seed were then exposed to control (18-20°C) or high temperature conditions (36°C, 38°C, 40°C, or 42°C) in a benchtop incubator (25 L, Vevor). The control and one elevated temperature were run each day with elevated temperatures run in random order (n=1 plate per temperature). Each plate contained n=84 seed and n=12 blank samples and were measured hourly for 4 h. After the final measurement, we assessed survival of all seed by rinsing in DI water and examining under a dissecting microscope. Seed were considered alive if the shell was closed or moved in response to probing or were considered dead if the shell was open and did not respond to probing.

Metabolism was calculated as described in the standard protocol. The effect of temperature on metabolism (size-normalized fluorescence; cube-root transformed) over time was tested using a linear mixed effect model (LMM) with hour and temperature as main effects, sample nested within plate within date, tank, and broodstock group as random intercepts in the *lme4* package (Bates et al., 2014) in R Statistical Programming v4.4.1 (R Core Team, 2022). Total metabolism over the trial was calculated as AUC and analyzed using linear mixed effect models with temperature as the main effect and tank and broodstock group as random effects. We then examined differences in metabolism between seed that were found to either be alive or dead at the end of the trials (at 42°C only as the other treatments did not show mortality) using a linear mixed effect model with time point and final mortality status as main effects with tank and broodstock group as random intercepts. Mortality effects on AUC were analyzed using linear mixed effect models with final mortality status as main effect with tank and broodstock group as random intercepts. For all models, significance was assessed using Type III Analysis of Variance tests with Satterthwaite’s approximation in the *lmerTest* package (Kuznetsova et al., 2015). Post hoc comparisons were evaluated using estimated marginal means tests in the *emmeans* package (Lenth, 2018). Temperature sensitivity (Q_10_) values were calculated as group means between 20°C and 36°C as well as between 36°C and 42°C using the following equation: (AUC at temperature 2 / AUC at temperature 1)^(10 / (temperature 2-temperature 1)) (Casas et al., 2018).

#### (2) Thermal response in medium seed (5-15 mm)

Oyster metabolism was measured across a range of temperatures to generate a thermal response curve for seed (5-15 mm) at the USDA Pacific Shellfish Research Unit in Newport, OR. Six temperatures were tested: 21°C, 27°C, 32°C, 37°C, 42°C and 45°C. Pediveliger oyster larvae were obtained from Whiskey Creek Hatchery (Tillamook, OR) and set with epinephrine at the Hatfield Marine Science Center (Newport, OR) to produce seed for the temperature trials. Seed were held in ambient sea water (12-16°C, 28-32 psu) at the Hatfield Marine Science Center until resazurin assays were conducted. For the experiments, oyster seed were placed individually into 12-well tissue culture plates (CellTreat) and photographed to measure size in ImageJ (Rueden et al., 2017) before adding 4 mL of resazurin working solution to each well. After resazurin working solution was added, 300 µL were sampled immediately from each well for an initial fluorescence measurement on a 96-well plate with fluorescence read on a Biotek Synergy LX plate reader (Agilent, Santa Clara, CA) with an emission of 528 nm (528/20 nm filter) and excitation of 590 nm (590/20 nm) with the Gen 5 software (Agilent, Santa Clara, CA). Well plates with seed and resazurin were then placed in a benchtop incubator set to one of the six temperatures with subsequent samplings and fluorescence readings of 300 µL of resazurin working solution every hour for 5 h with n=5 blanks per trial. Trials were performed over a series of 6 days with one temperature run each day in random order. Mortality was not assessed in this experiment.

Data were analyzed as described above. The effect of temperature on metabolism was tested with two-way ANOVA models with temperature and time point as the main effects. The effect of temperature on AUC was tested by using a one-way ANOVA model with temperature as the main effect. Q_10_ values were calculated as group means between each pair of temperatures (e.g., 21°C to 27°C, 27°C to 32°C, and so on) as described above.

### D. Objective 3: Characterization of acute thermal stress responses

We examined oyster metabolic response to acute temperature stress by exposing seed (15-35 mm) to either control (18°C; n=20 oysters per trial) or high (42°C; n=20 oysters per trial) temperatures for 4 h. Oyster seed were obtained from the Point Whitney Jamestown S’Klallam Shellfish Hatchery and held at 14°C in tanks filled with water adjusted to 25 psu using Instant Ocean salts (Instant Ocean, Blacksburg, VA) at the University of Washington. Seed were obtained from a separate experiment in which seed were exposed 2 months prior to weekly 1 h exposures to temperature (25°C), fresh water, or immune (Poly(I:C) exposure) sublethal stressors (n=2 replicate bags of oysters per treatment). Treatment effects were not significant on metabolism and are therefore included as a random effect but are not evaluated here directly (see Results). Seed were imaged for shell length (mm) and placed in plastic cups with lids filled with 20 mL of resazurin working solution made as described above. Fluorescence measurements were taken by sampling 200 µL of the resazurin solution from each cup, placing in a 96-well plate, and reading fluorescence on a plate reader (PerkinElmer Victor 3, Shelton, CT) in fluorescence mode with emission of 528 nm (528/20 nm filter) and excitation of 590 nm (590/20 nm filter) with associated software (PerkinElmer Victor X Software). Due to the larger size of seed, we were able to visually assess the survival of each oyster every hour while seed remained in plastic cups. Seed were considered alive if the shell was closed or moved in response to probing or were considered dead if the shell was open and did not respond to probing. Trials (n=10) were repeated over ten days (N=506 oysters) with n=6 blanks per trial.

Metabolism was calculated as described above. We analyzed the effect of temperature on oyster metabolism that survived the trial and the effect of mortality status on the metabolism between those that either lived or died during the trial at 42°C. For each analysis, metabolism over time was tested using a linear mixed effect model with hour and temperature or mortality as main effects, sample nested within bag and date for repeated measures, and prior stress treatment as random intercepts in the *lme4* package (Bates et al., 2014). Effects of prior stress treatment were evaluated using ANOVA-like tests for random effects in the *lmerTest* package (Kuznetsova et al., 2015). We also analyzed the effect of temperature or mortality on AUC with bag and date as random intercepts. For all models, significance was assessed using Type III Analysis of Variance tests with Satterthwaite’s approximation in the *lmerTest* package (Kuznetsova et al., 2015) and post hoc tests were conducted using estimated marginal means tests in the *emmeans* package (Lenth, 2018). Survival was analyzed using a logistic regression (GLMM; *glmer* function; family=binomial, link=logit) in the *lme4* package (Bates et al., 2014) with binary mortality as the response with time and temperature as main effects and prior stress treatment, bag, and sample (for repeated measures) as random intercepts. Significance was assessed using Type II Wald chi-square ANOVA tests in the *car* package (Fox and Weisberg, 2018).

Finally, we evaluated whether metabolism at a particular time point predicted mortality at the subsequent time point. We assessed the relationship between metabolism and subsequent mortality using a logistic regression model (GLMM; *glmer* function; family = binomial, link = logit) in the *lme4* package (Bates et al., 2014). Mortality at the subsequent time point was included as the response variable with scaled fluorescence and temperature as main effects. Prior stress treatment and sample nested within bag and date were included as random intercepts. Model significance was evaluated using a Type II Wald chi-square ANOVA test in the *lmerTest* package (Kuznetsova et al., 2015). Predicted probability of mortality was then plotted against metabolism with confidence intervals generated using the *pROC* package (Robin et al., 2011).

### E. Objective 4: Examination of genetic variability in metabolic responses

To examine genetic variation in metabolism between oyster families, we obtained oyster seed (13-25 mm) from the United States Department of Agriculture Pacific Shellfish Research Unit Pacific Oyster Genome Selection program (Newport, OR) from five families produced during the 2024 spawning season. Oysters were held in heath trays at ambient conditions at the Jamestown S’Klallam Shellfish Hatchery (Brinnon, WA) until April 2025 at which point they were transported to the University of Washington. Oysters were then held at 14°C in tanks filled with water adjusted to 25 psu using Instant Ocean salts (Instant Ocean, Blacksburg, VA) at the University of Washington.

Resazurin solutions were prepared as described above and added to oysters placed in 20 mL plastic cups. Fluorescence readings were taken by sampling 200 µL of the resazurin solution and reading on 96-well plates on a BioTek Synergy HTX multi-mode plate reader (Agilent, Santa Clara, CA) in fluorescence mode with emission at 530 nm (528/20 nm filter) and excitation at 590 nm (590/20 nm filter) with associated software (Agilent Gen5 Software). Following initial measurement, all oysters were placed in an incubator set to the control temperature (18°C) and samples were collected every hour for 3 h. After the 3 h sample was taken, the incubator was set to 40°C and oysters were returned to the incubator with samples taken every hour for 3 h. The total incubation time was 6 h with the first half at control temperature and the second half at high temperature. This approach allows for tracking individual oyster metabolic response to acute temperature stress. Temperatures in oyster cups were monitored using a digital thermometer (Traceable Excursion-Trac, San Francisco, CA). Following the final measurement at 6 h, oysters were assessed for mortality as described above with analysis conducted only for oysters that survived. Each trial included n=5 oysters per family and n=5 blank samples with trials (n=6) conducted over 6 days (N=150 oysters, n=30 per family).

Metabolism calculations were performed for each individual during control and elevated temperature phases. The effect of family and temperature on metabolism (cube-root transformed) over time was tested using a linear mixed effect model with family, time point, and temperature as main effects, sample nested within date (for repeated measures), and measurement date as random intercepts in the *lme4* package (Bates et al., 2014) in R Statistical Programming v4.4.1 (R Core Team, 2022). For all models, significance was assessed using Type III Analysis of Variance tests with Satterthwaite’s approximation in the *lmerTest* package (Kuznetsova et al., 2015) and post hoc tests were conducted using estimated marginal means tests in the *emmeans* package (Lenth, 2018).

### F. Objective 5: Conduct a case study to identify relationships between metabolism and performance in *Crassostrea virginica*

#### (1) Selective breeding program

We conducted resazurin assays on selectively bred families to test relationships to predicted performance at the Virginia Institute of Marine Science (VIMS). The Aquaculture Genetics and Breeding Technology Center (ABC) at VIMS has been performing family-based breeding of *Crassostrea virginica* since 2004. A full description of the diploid family breeding program at ABC is outlined in (Allen et al., 2021). Current traits for selective breeding include survival, total weight, meat yield and shell shape characteristics across low and moderate/high salinity environments, with a strong genetics x environment effect for survival and growth traits between low and moderate/high salinity sites. Note that moderate/high salinity test sites often have annual disease pressure from *Perkinsus marinus* and *Haplosporidium nelsoni*, both of which are linked to oyster mortality. Phenotypic information is collected on families at 18 months of age and selection of broodstock candidates for subsequent spawns and terminal broodstock line creation is based on two selection indexes, one for low salinity performance and one for moderate/high salinity performance. Selection locations have varied over the years; however, current selection locations include those in **Table S1**. Resazurin assays in this study were conducted using thermal stress tests of metabolism as an indicator of stress tolerance to assess whether metabolism corresponds to general performance.

#### (2) Resazurin assays on selectively bred families

For this study, all seed were spawned in spring 2025 at the Acuff Center for Aquaculture at VIMS by members of ABC. Resazurin assays were performed on *C. virginica* oysters from 50 families at VIMS with a total of 28 oysters per family measured (n=1,400 oysters). Family spawns were performed using 1x1 crosses of broodstock candidates selected for specific performance traits (described below) where each cross was kept separate through the entire larval and setting period. Larvae for family crosses were reared in 60 L static, aerated tanks whereas larvae for lines were reared in 200 L tanks. Larval husbandry was generally the same for all cultures, including daily feeding of live microalgae cultured at the Acuff Center, water changes, tank cleaning and larval health assessments every other day. When the larvae approached 300 µm in size and developed an eye spot, they were added into coffee filters dampened with seawater and removed from each tank. Harvests were stored at 4°C and occurred every other day for 3 days. Harvests were then combined and eyed larvae were set on 400 µm microcultch in the downwelling setting system within the Acuff Center. While in the setting system, setters were fed daily a ration of live microalgae, and each setting tank was cleaned every other day. Once the setters reached ∼1mm in size, the cultch was removed through sieving, and all seed were transferred to the land-based upweller system in the VIMS boat basin. One day before the resazurin assays, a subset of seed from each family was removed from each silo and set aside by ABC and placed into an individually labeled mesh pouch. Pouches were then held briefly in three upweller silos before being transferred to the Acuff Center broodstock room. Seed used for resazurin measurements ranged from 7-25 mm in size.

Families were randomly chosen to include those selectively bred for either moderate/high salinity (n=25 families) and low salinity (n=25 families) performance. Assays were conducted in 24-well plates (2.5 mL) with n=4 blanks and n=20 oysters per plate. Prior to assays, we conducted preliminary observations to select temperature treatments that maximized stress exposure but did not cause mortality. We selected 40°C for 2.5 h followed by a recovery period for another 1.5 h as our case study trial. Oysters were randomly allocated to plates for a total of n=70 plates with n=22-24 plates run each day over three days. Resazurin solutions were prepared as described and measurements were collected on a Varioskan Lux (Thermo Scientific, Waltham, MA, USA) multi-mode plate reader at an excitation at 530 nm and emission at 590 nm with a bandpass of 12 nm using SkanIt software. Chilled (4°C) resazurin was added to plates for initial measurements and plates were then added to an incubator set at 40°C. Plates were added to incubators at staggered intervals to ensure each plate was read every hour. Plates were measured again at 1 and 2 h time points. At 2.5 h, plates were moved out of the incubator for a recovery phase. Measurements were then collected at 3 and 4 h time points. Temperature was recorded in one randomly selected well in 12-16 plates hourly. 145 samples (10%) reached oversaturation and were removed from the analysis. Temperature profiles of the wells are displayed in **Fig S1.** Temperatures were 12.3±0.3°C at initial measurements, 34.2±0.1°C at 1 h, 35.6±0.2°C at 2 h, 29.0±0.3°C at 3 h, and 23.2±0.2°C at the final measurement at 4 h. We assessed each individual for mortality at the end of the trial.

#### (3) Data analysis and correlation to predicted performance

Metabolism was calculated as described previously for each oyster. We analyzed metabolism over time using a linear mixed effect model with cube-root transformed fluorescence as the response and family and time point and their interaction as main effects using the *lme4* package in R (Bates et al., 2014). Well, plate, and date were included as random intercepts to account for repeated measures. We also tested for the effect of family phenotype (i.e., low or high salinity selected) using mixed effect models with the same structure but with time point and phenotype as main effects. Significance of main effects was evaluated using Type III ANOVA tests with Satterthwaite’s approximation method. Normality of residuals and homogeneity of variance were assessed using quantile-quantile plots and Levene’s tests, respectively. AUC was analyzed using a linear mixed effect model with family or phenotype as the main effect with plate and date as random intercepts. We used a repeatability estimation in the *rptR* package (Stoffel et al., 2017) to evaluate intraclass correlation and describe within and between-group sources of variation. Family, as the grouping variable, was included as a random intercept with a main effect of time point and fold change in fluorescence as the response with Gaussian data.

Predicted performance data after progeny testing is analyzed annually using linear mixed animal models using a genetic groups structure in AsReml (Gilmour et al., 2015) as in (Allen et al., 2021). Growth traits are analyzed using a five-trait, multivariate model whereas survival is analyzed in a separate model. “Predicted performance” values were calculated for each family as estimated breeding values for survival and growth in low or moderate/high salinity environments (percentage gain) generating four metrics: predicted survival in moderate/high salinity, predicted survival in low salinity, predicted growth in moderate/high salinity, and predicted growth in low salinity environments. We then correlated AUC of each oyster with the predicted performance metrics for survival and growth using non-parametric Spearman correlation tests without data transformations. We conducted correlation analyses between metabolism and predicted performance metrics for high and low salinity selected families separately.

## Results

### A. Objective 1: Adaptation of resazurin assay to measure whole organism oyster metabolism

#### (1) Positive correlation between resazurin and oxygen measurements of metabolism

We first explored the relationship between metabolism measurements made through resazurin fluorescence measurements and oxygen consumption. There was a significant positive correlation between oxygen consumption and resazurin fluorescence (Pearson correlation; p=0.012, r=0.321; **Fig S2**). This correlation was weakly positive with r=0.321.

#### (2) Clear fluorescence signals from live animals

We compared fluorescence signals from blank, empty shells, and live oysters to confirm that metabolism of live animals is not confounded by signals originating from microbial communities on oyster shells or reactions between the shell material and resazurin solution. We found that signals from empty shells were higher than blanks (ANOVA; SS=14.4, F=111.4, p<0.001; **Fig S3**), but that live oyster metabolism was dramatically higher than empty shells (ANOVA; SS=4.9, F=96.4, p<0.001; **Fig S3**).

#### (3) Metabolism correlates positively with oyster size

Total metabolism over the course of the trials (calculated as AUC) positively correlated with oyster size (Spearman correlation, S=2.82e6, p<0.001) with a moderately positive relationship (r=0.624; **Fig S4**). Length effects on AUC were significant (LM; t=13.90, adj. R^2^=0.13, p<0.001) with a slope of 0.13, indicating a one unit increase in AUC with every 0.13 mm increase in length.

### B. Objective 2: Strong temperature effects on metabolism

Metabolism was strongly influenced by temperature. We examined temperature effects on metabolism using the resazurin assay in small seed (3-8 mm) measured at the University of Washington as well as seed (5-15 mm) measured at the USDA Pacific Shellfish Research Unit (Newport, OR). Metabolism in small seed (3-8 mm), including only those that survived the trial, varied across temperatures (LMM; SS=3.05, DF=16/2260, F=15.49, p[time point x temperature]<0.001). Total metabolism over the course of the incubation, measured as area under the curve (AUC), changed with temperature (LMM; SS=5.28, DF=4/562.5, F=22.36, p<0.001) with rates peaking at 36°C and decreasing at 38-42°C (**Fig 2A**). Q_10_ values were highest from 20-36°C (Q_10_=1.22) followed by decreased thermal sensitivity from 36-42°C (Q_10_=0.28; **Table S2**). Similarly, in larger seed (5-15 mm) metabolism differed across temperatures (LMM; SS=4.91, DF=24/625.53, F=37.24, p[time point x temperature]<0.001). AUC varied strongly across temperatures (LMM; SS=7.29, DF=5, F=32.85, p<0.001) with metabolic activity increasing from 21-26°C to a peak at 37-42°C and declining sharply at 45°C (post hoc p<0.05; **Fig 2B**). Thermal sensitivity (Q_10_) peaked between 26-32°C (3.52) and was lowest in temperatures exceeding the thermal optimum between 32-45°C (Q_10_=0.001-1.22; **Table S2**).

**Fig 2.**
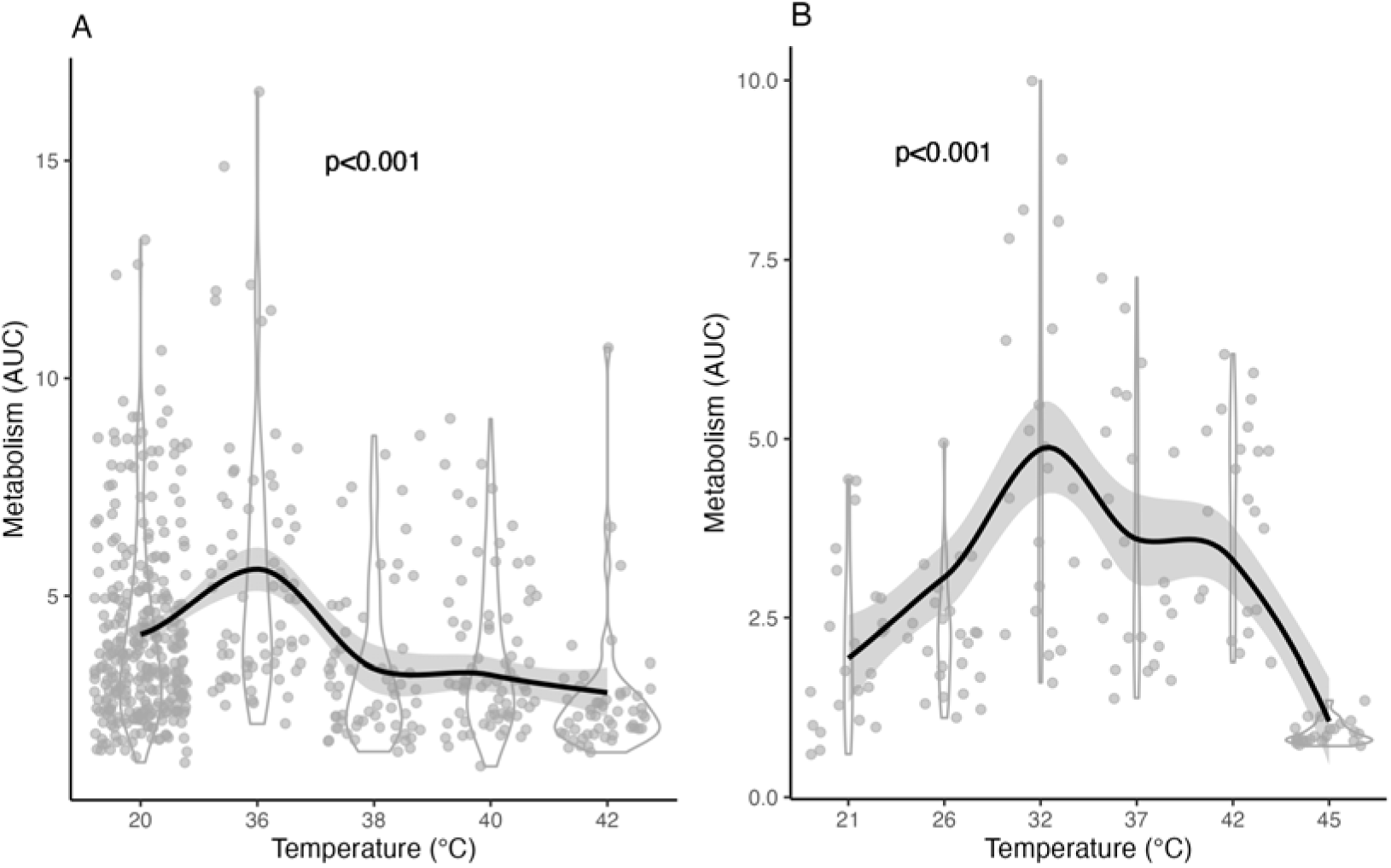
(A) Metabolism (area under the curve; AUC) across temperatures (°C) for small seed (3-8 mm) analyzed at the University of Washington. Plot displays only seed that survived the trial. (B) AUC across temperatures for seed (5-15 mm) analyzed at the Pacific Shellfish Research Unit. In all plots, loess line displays response across temperatures.

### C. Objective 3: Clear metabolic response to acute stress and metabolic depression as an indicator of tolerance

We observed strong metabolic responses to acute high temperature stress using the resazurin assay and identified metabolic indicators of resilience. In large seed (15-35 mm) that survived the entire 4 h incubation (n=92 at 42°C, n=201 at 18°C), metabolism was significantly different between 18°C and 42°C (LMM; SS=1.28, DF=4/11184, F=116.62, p[time point x temperature]<0.001) with oysters exposed to 42°C exhibiting metabolic depression (**Fig 3A**). Metabolism was significantly lower at 42°C from 2-4 h of incubation (post hoc p<0.001). Total metabolism (calculated as AUC) in oysters that lived during the 4 h incubation was lower at 42°C than 18°C (LMM; SS=1.54, DF=1/291, F=63.39, p<0.001; **Fig 3B**). Note that previous stress exposure had no effect on metabolic rates (ANOVA-like test for random effects; log likelihood=1905.1, likelihood ratio test=0.00, DF=1, p=1).

**Fig 3.**
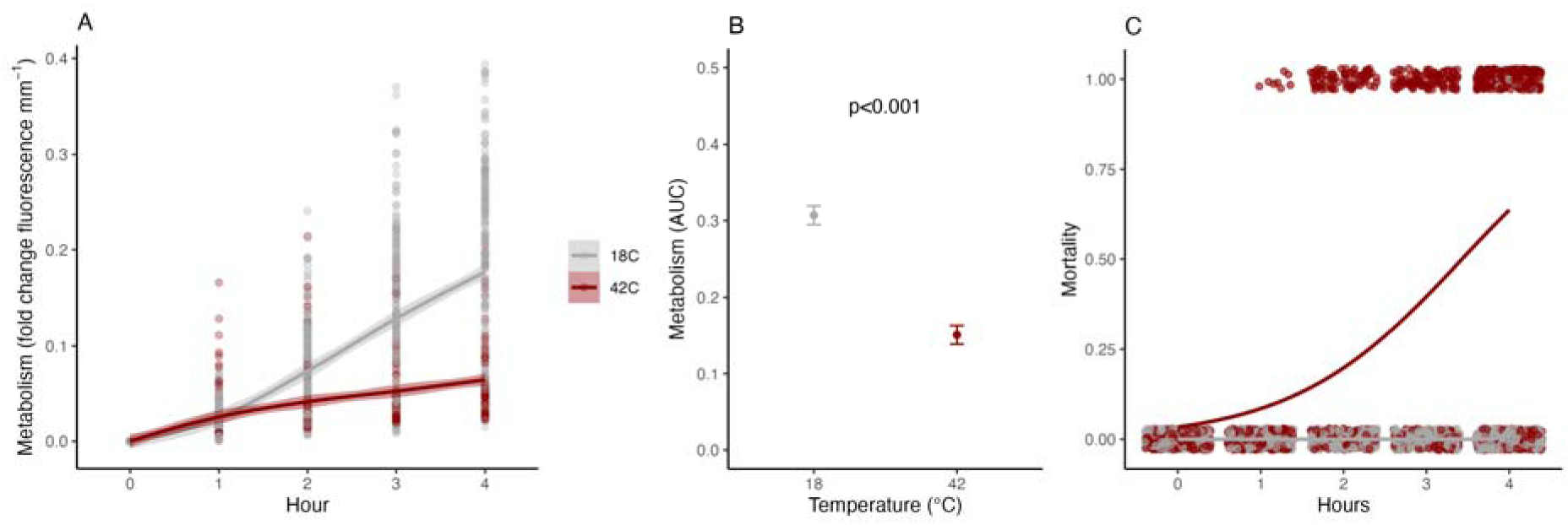
(A) Metabolism (fold change in fluorescence mm^-1^ over time) in oysters that survived the 4 h incubation at 18°C (gray) or 42°C (red). (B) Metabolism (area under the curve, AUC) at 18°C and 42°C. (C) Mortality during the 4 h incubation period. Lines indicate binomial mixed effect model predications with a value of 1 indicating mortality and a value of 0 indicating a live oyster.

Mortality was higher at 42°C than 18°C during the 4 h resazurin measurement (GLMM; X^2^=115.31, DF=1, p<0.001; **Fig 3C**). During the trials, 58±3% of oysters died at 42°C while only 4±1% died at 18°C. Given the degree of mortality at 42°C, we then examined differences in metabolism between oysters that lived (n=92) and those that died (n=127) during the 4 h exposure at 42°C.

First, we examined metabolism in oysters depending on time of mortality and observed that oysters that died earlier in the trial (1-3 h) showed slower metabolic rates than those that died at the end of the trial (4 h) (**Fig S5**). Further, oysters that remained alive showed continued increases in metabolism throughout the 4 h measurements, in contrast to reduced metabolism in those that died (**Fig S5**). Overall, metabolism was lower in oysters that lived compared to those that died during the 4 h exposure to 42°C. Metabolism was lower in large seed (15-35 mm) that lived compared to those that died during the 4 h measurements (LMM; SS=0.51, DF=4/849.95, F=133.19, p[time point x mortality]<0.001; **Fig 4A**) as was AUC (LMM; SS=1.26, DF=1/212.58, F=95.791, p<0.001; **Fig 4B**). Metabolism was lower in oysters that survived (post hoc p<0.001; **Fig 4A**) with a 52% decrease in AUC in oysters that survived (post hoc p<0.001; **Fig 4B**). Similarly, in small seed (3-8 mm), metabolic rates (LMM; SS=0.30, DF=4/404, F=5.15, p[time point x mortality]<0.001) and AUC (LMM; SS=3.0, DF=82, F=14.71, p<0.001) were lower in oysters that survived the 42°C exposure than those that died during the incubation (**Fig 4CD**).

**Fig 4.**
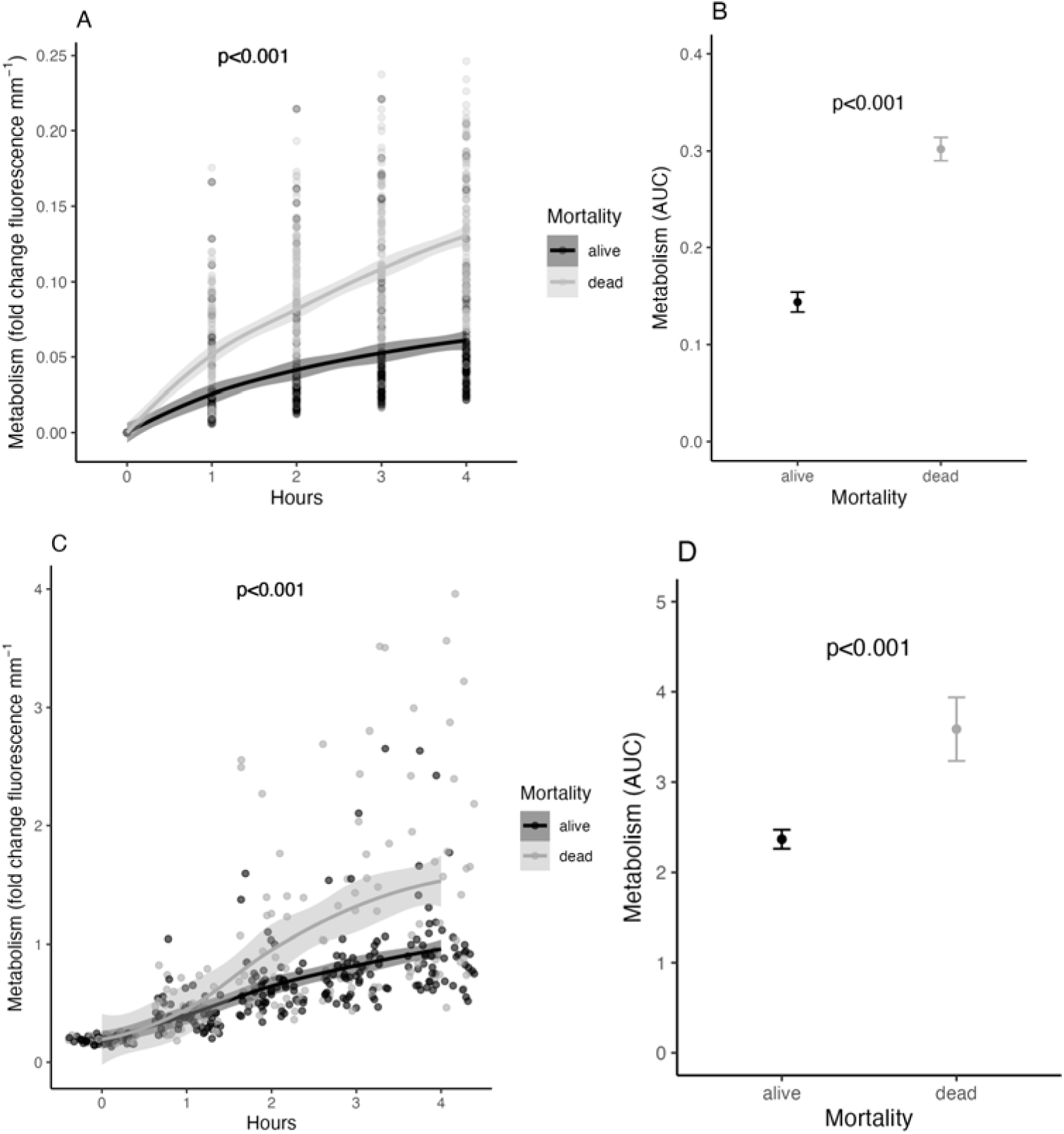
(A) Metabolism (fold change in fluorescence mm^-1^ over time) in oysters (15-35 mm) that survived (black) or died (gray) by the end of the trial at 42°C. (B) Total metabolism (area under the curve, AUC) in oysters that survived (black) or died (gray) by the end of the trial at 42°C. (C) Metabolism (fold change in fluorescence mm^-1^ over time) in oyster seed (3-8 mm) that survived (black) or died (gray) by the end of the trial at 42°C. (D) Total metabolism (area under the curve, AUC) in oyster seed (3-8 mm) that survived (black) or died (gray) by the end of the trial at 42°C.

We further evaluated whether metabolism at a particular time point predicted mortality at the subsequent time point under acute stress. Higher metabolism was significantly associated with increased odds of mortality (GLMM; estimate =4.0, X^2^=67.09, DF=1, p<0.001; **Fig 5**). In other words, oysters with high metabolism are at greater risk for mortality (**Fig 5**).

**Fig 5.**
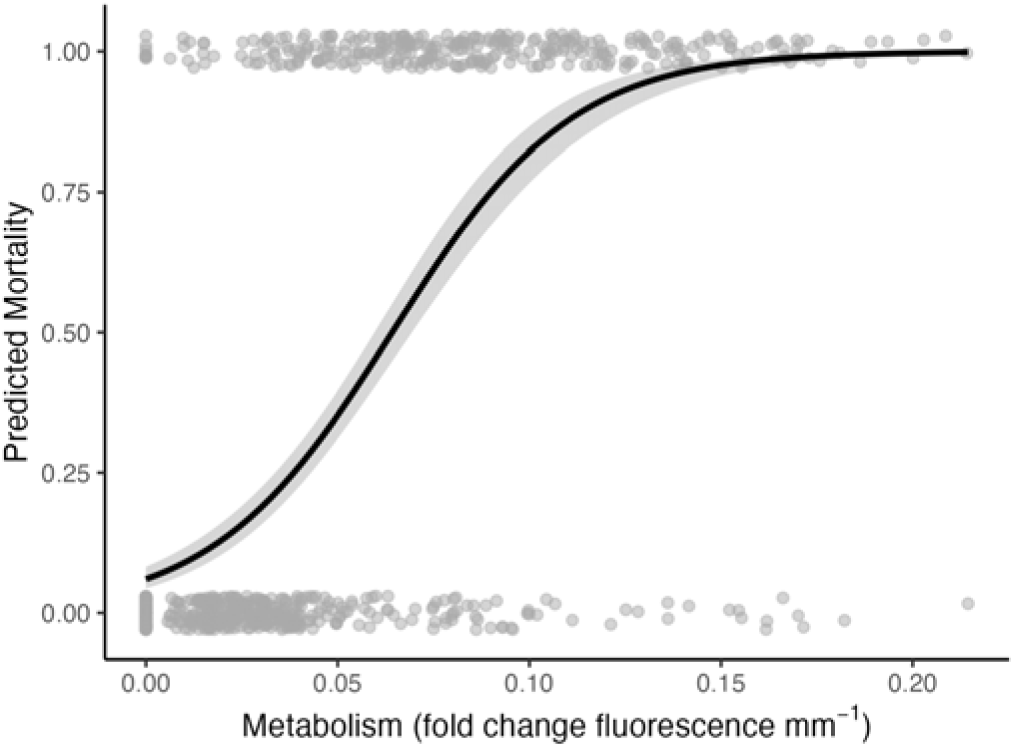
Predicted mortality associated with metabolism (fold change in fluorescence mm^-1^) measured using the resazurin assay at 42°C. Line indicates model prediction of the relationship between metabolism and mortality. Data points indicate observations in which 0 indicates a live oyster and 1 indicates a dead oyster by the end of the trial.

### D. Objective 4: Genetic variation in metabolism response

We conducted resazurin measurements on five families of oyster seed (13-25 mm) exposing each individual to 3 h at 18°C followed by 3 h at 40°C to examine genetic variation in metabolism. Responses varied between families (LMM; SS=0.05, DF=4/934.37, F=3.45, p=0.008) and metabolic rates were higher at 40°C than at 18°C (LMM; SS=0.02, DF=1/923.53, F=6.09, p=0.014; **Fig 6A**), but there were no significant interactive effects between temperature and family (LMM; SS=0.01, DF=4/926.87, F=0.44, p=0.782; **Fig 6B**). Within each family, there was variation between individuals, but metabolism was higher in families A and B than in families C and D (Fig **6A****B**).

**Fig 6.**
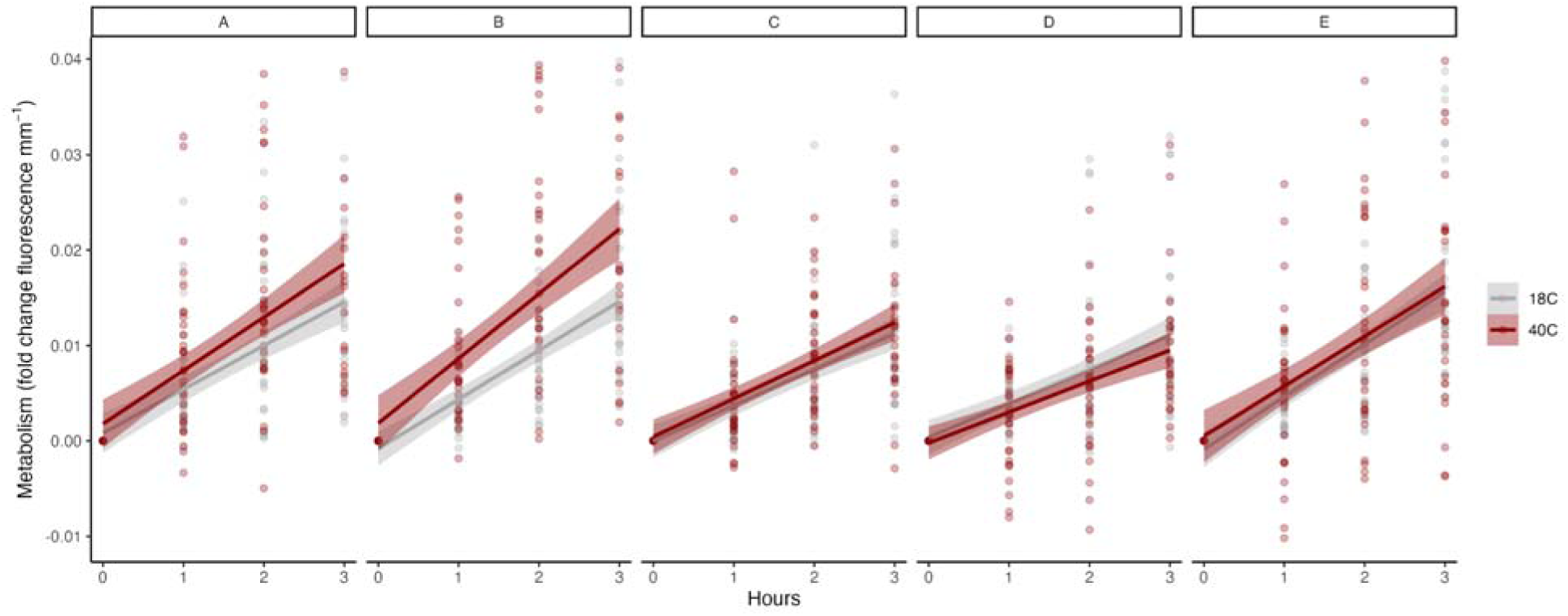
(A) Metabolism (fold change in fluorescence mm^-1^) for each oyster family (families A-E) exposed to 18°C for 3 h (gray) followed by 40°C (red) for another 3 h. Lines indicate mean response in each temperature treatment.

### E. Case study: Metabolism correlates with predicted performance in *Crassostrea virginica*

We applied the resazurin metabolic assay in a case study of *C. virginica* seed (7-24 mm) to examine whether metabolism under heat stress correlates with predicted performance in 50 selectively bred families. All oysters were exposed to 2.5 h of acute heat stress followed by a recovery period (**Fig S1**). There was significant variation in metabolism between families (LMM; SS=30.74, DF=196/4788, F=4.56, p[family x time]<0.001; **Fig 7A**). There was also a significant effect of family selected phenotype (i.e., high/moderate salinity or low salinity selected families) on metabolic rates (LMM; SS=1.29, DF=4/4980, F=8.24, p[phenotype x time]<0.001; **Fig 7B**). Similarly, total metabolism over the course of the trials (AUC) varied significantly between families (LMM; SS=136.83, DF=49/1146, F=5.02, p<0.001; **Fig 7C**) and between selected phenotypes (LMM; SS=5.60, DF=1/1239.8, F=8.77, p=0.003; **Fig 7D**). Specifically, metabolism was higher in the low salinity selected families. Repeatability analyses indicated that most variation in metabolism was due to individual response within families with little structure between families (R=0.11, p<0.001).

**Fig 7.**
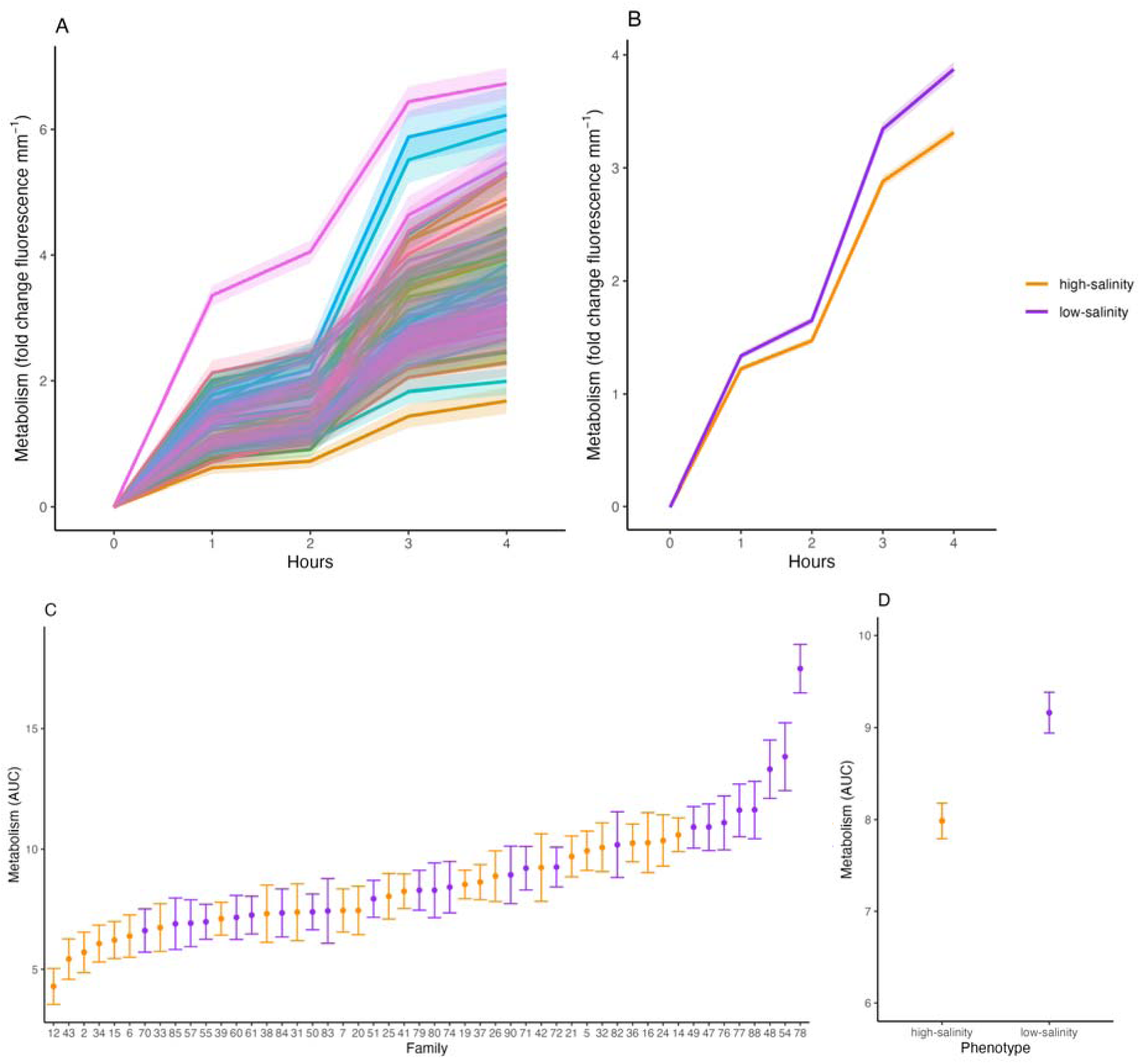
(A) Metabolism (fold change in fluorescence mm^-1^) of each family (represented by color) during resazurin measurements. (B) Metabolism of each selected phenotype (i.e., selected for high/moderate salinity or low salinity performance). A-B, lines indicate the modeled response with error bars indicating 95% confidence intervals. (C) Total metabolism (area under the curve; AUC) for each family and (D) AUC summarized by selected phenotype. In B-D, families selected for high/moderate salinity environments are in orange and those selected for low salinity environments are in purple.

We then correlated mean AUC of each family with the family’s predicted field performance metric. We tested for correlations between AUC and growth and survival in each phenotype group (high/moderate salinity or low salinity selected). We found that AUC in low salinity selected families was significantly positively correlated with predicted survival in a low salinity environment (Pearson correlation; t=3.16, df=22, r=0.56, p=0.004; **Fig 8**). However, there was no significant correlation within the high salinity selected families, but there was a trend for a negative relationship (Pearson correlation; t=-1.38, df=22, r=-0.28, p=0.181; **Fig 8**). Interestingly, there was no significant correlation between AUC and predicted survival in high salinity environments (Pearson correlation; p>0.05). There was no significant correlation with predicted growth for any phenotype group (Pearson correlation; p>0.05). There was no mortality observed during or after the resazurin trials.

**Fig 8.**
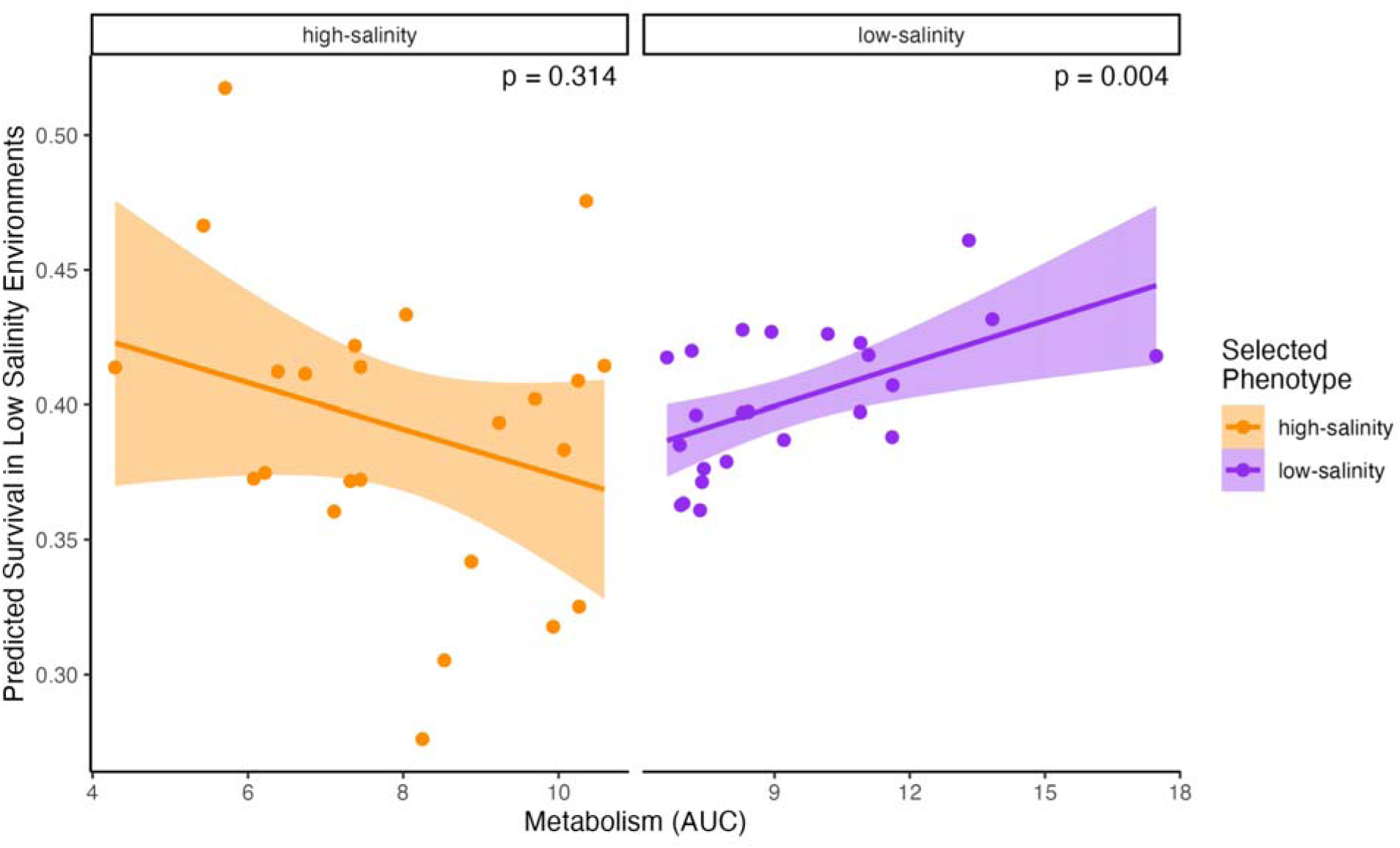
Correlation between total metabolic activity (AUC) and predicted survival (% gain) in low salinity environments. Correlations conducted separately for families selected for performance in low salinity (purple) and high/moderate salinity (orange) environments.

## Discussion

This study establishes resazurin metabolic assays as a reliable tool for measuring oyster metabolism, with clear utility for both scientific research and aquaculture applications. By adapting a dye widely used in a variety of cell viability, microbial, and biomedical assays (e.g., Anoopkumar-Dukie et al., 2005; Petiti et al., 2024; Zare et al., 2015) and more recently applied in fish (Reid et al., 2018; Renquist et al., 2013), we demonstrate that resazurin provides a means of quantifying whole-organism metabolism in oysters. Quantifying metabolic responses is key to understanding health, performance, and survival in challenging and energy-limiting environments (Sokolova, 2013). Our experiments show that this approach can provide a high-throughput and reliable method to capture variation in thermal performance, acute stress responses, and family-level differences in metabolism. Our application in an aquaculture case study further demonstrates that metabolism can be correlated to predicted performance in oysters, providing a foundation from which future work can identify linkages between metabolism and field performance. The adaptation of this assay for use in shellfish opens a wide range of new opportunities to further optimize and develop standard operating procedures for this approach to measure performance, monitor stress, identify resilient stocks, investigate physiological responses of shellfish for scientific and aquaculture applications.

Our first goal was to validate the use of resazurin assays in reliably measuring metabolism in oysters. Resazurin assays provide a robust measure of metabolism by acting as an intermediate electron acceptor and changing from the oxidized, non-fluorescent form (blue) to the fluorescent resorufin (pink) form (Rampersad, 2012). As the organism conducts metabolic activity the dye is reduced by NADH in the presence of reductases and fluoresces (Candeias et al., 1998; Chen et al., 2018; O’Brien et al., 2000), which is measured using a plate reader. Our results confirm that the resazurin assay measures metabolic signals as biologically expected from live animals (i.e., metabolism was far higher in live animals than from empty shells), that signals increase with animal size, and that metabolism measured via resazurin is positively correlated with oxygen consumption. These tests further highlight differences in measurement approaches between resazurin fluorescence and other widely used methods such as oxygen consumption. The relationship between oxygen consumption and resazurin fluorescence was positive, but weak, demonstrating variability between the two measurements. This may reflect differences in metabolic pathways measured (e.g., aerobic *versus* anaerobic processes), underscoring that resazurin provides complementary, rather than redundant, information about metabolism compared to oxygen consumption approaches. Variability in measurements between the two approaches is likely driven by individual oyster day-to-day variation in responses as well as the difference in mechanism of measurement of respiration (i.e., oxygen consumption) *versus* resazurin (i.e., electron transport). Under temperature stress, *Crassostrea gigas* oysters exhibit transitions from aerobic to anaerobic metabolism (Li et al., 2017), suggesting a need for metabolic assays that capture multiple metabolic pathways and metabolic plasticity. Indeed, resazurin assays may capture a wider range of metabolic responses than anaerobic metabolism alone and can be adapted to capture metabolic activity in hypoxic conditions (Lavogina et al., 2022).

One promising application of the resazurin assay is the rapid and high throughput assessment of metabolic performance across environmental performance curves (e.g., thermal performance curves). In our results, we observed peak metabolism at 32-36°C followed by metabolic depression at higher temperatures in multiple experiments. Similar patterns have been observed in previous studies in oysters. For example, observations of heart rate in *Ostrea edulis* show peak cardiac activity 30-34°C followed by decreases near the lethal temperature of 34-36°C (Eymann et al., 2020; Götze et al., 2025). Similarly, Tropical *Isognomon nucleus* oysters exposed to increasing temperatures showed peak oxygen consumption at 39-41°C, depending on thermal history, after which oxygen consumption declined (Giomi et al., 2016). Respiration rates in *Crassostrea gigas* also display increased metabolic activity at 35°C compared to 20°C (Li et al., 2017). Further, temperature sensitivity of metabolism, calculated as Q_10_, ranged from <1 to ∼3.5 in our study, which are similar to observations in respiration of oysters in previous studies (e.g., Q_10_ values of ∼1.4 (Pan et al., 2021) and ∼1.3-2.2 (Casas et al., 2018)). Our observation of expected temperature-dependent metabolic activity using the resazurin assay demonstrates the utility of this approach in characterizing thermal performance curves in oysters at high replication. However, we did not conduct oxygen saturation measurements during trials and even though plates were unsealed, allowing for oxygen diffusion into the resazurin solution, it is possible that thermal stress effects were combined with oxygen stress. We recommend that trials include oxygen saturation measurements to confirm oxygen saturation in solution.

Importantly, the resazurin assay detected metabolic signatures associated with survival under acute stress conditions in our study. In acute stress tests, oysters that survived exhibited stronger metabolic depression compared to those that died, indicating that metabolic downregulation may serve as a protective mechanism. Metabolic depression may serve as a strategy to conserve resources and allow for survival in the short term (Pörtner, 2008; Sokolova, 2021) but if prolonged, it becomes unsustainable with a risk for negative consequences (Sokolova, 2013). In intertidal tropical oysters (*Isognomon nucleus*), metabolic depression may play a role in energy conservation during periods of short-term heat stress (triggered at 37°C) at low tides (Hui et al., 2020). Oysters also exhibit metabolic depression in response to environmental stressors including high pCO_2_/acidification conditions (Lannig et al., 2010; Le Moullac et al., 2007) and starvation (García-Esquivel et al., 2002). We hypothesize that under short term acute stress tests, oysters with greater capacity for metabolic depression may represent more resilient stocks to acute stress events and our results suggest that short-term resazurin assays could provide a rapid means of assessing stock resilience to thermal stress. Future applications of acute stress tests using the resazurin assay must first consider local environmental context and conduct preliminary testing to determine acute temperatures which may elicit metabolic depression as a survival response, without complete mortality. It is important to note that even after a behavioral determination of mortality there is still residual metabolic activity detected using the resazurin assay due to ongoing metabolic activity in oyster tissues and microbial communities. Therefore, any determinations of mortality made using resazurin assays should be complemented and confirmed by behavioral and observational determinations of survival. While responses will need to be contextualized for each system and species, resazurin assays are a potential option for rapid stress-testing in hatcheries and monitoring programs. A critical next step is to track and compare resazurin stress responses with field deployment of oyster stocks to determine if lab-based stress tests provide realistic estimates of performance in natural and aquaculture environments.

Genetic and family-level signatures in metabolic response highlight the potential application of resazurin in selective breeding programs and to provide an index of stock resilience. We observed variation in metabolism between oyster families, and in our case study with *C. virginica*, metabolism under temperature stress varied depending on the selective breeding phenotype. We utilized thermal stress testing as a rapid assay with resazurin to provide an indicator of oyster metabolic response under stress. Although the oysters in this case study were not bred specifically for thermal tolerance, the conserved energetic responses to stress provide an opportunity to relate single stressor response to generalized performance. Interestingly, metabolism was also correlated with predicted survival in low-salinity environments, but only for the families that were selectively bred for high performance in this environment. This finding may reflect shared physiological mechanisms between osmoregulation and thermal stress, as both require substantial energetic investment in maintaining cellular homeostasis (Sokolova, 2021, 2013) and the application of a thermal stress test assay of metabolism may provide an indicator of stress response that relates to capacity for generalized performance. In contrast, families bred for high-salinity environments are generally selected in environments with stronger disease and pathogenic pressure (Bushek et al., 2012; Haskin and Ford, 1982), where resilience may be shaped more by immune response (Frank-Lawale et al., 2014) than metabolic or osmoregulatory functions. These differences highlight that selective pressures targeting different stressors may favor distinct physiological pathways, and that metabolic assays may be useful for detecting matches and mismatches between selectively bred stocks and their potential environments. Further, we found that a small number of highest and lowest performing families drove the differences in metabolism between high and low salinity selected families, emphasizing the utility of the assay to detect high and low performers. These results also demonstrate the need to sample a high sample size and sample across multiple families to account for biological variability and allow for sufficient detection of high and low performers. With further validation against observed field performance, resazurin assays could provide hatchery managers with an accessible method to screen large numbers of individuals or families for resilience traits.

Overall, our work demonstrates that resazurin metabolic assays provide a valid and versatile approach for measuring metabolism in oysters and opens a wide range of opportunities to investigate oyster physiological responses to a variety of environmental conditions. Resazurin assays further offer a bridge between basic research and applied aquaculture, enabling scientists to probe physiological mechanisms of stress tolerance while providing growers and managers with an additional tool in aquaculture settings. Potential applications include screening for high-performing or stress-resilient stocks, monitoring health and stress in hatcheries, and characterizing family-level variation to inform breeding. It is critical that additional development of resazurin assays includes thorough testing and consideration of local environmental context to develop resilience or performance indices that are relevant for the stocks being tested. With further refinement and validation in field settings, the resazurin assay could become an important addition to the toolkit for oyster aquaculture and conservation under climate change.

## Acknowledgements

We would like to acknowledge and thank Jamestown S’Klallam Seafood - Point Whitney Hatchery, the United States Department of Agriculture Pacific Shellfish Research Unit Pacific Oyster Genome Selection program, and Pacific Hybreed for providing oysters used in this research. We acknowledge members of the Roberts Lab (University of Washington) for assistance with laboratory sampling and feedback on earlier drafts of this manuscript. We would also like to acknowledge Katie McFarland (NOAA) for her contributions to early development of the resazurin assay in eastern oysters. We thank Christopher P. Tatara (NOAA) for comments that improved the manuscript.

## Data Availability

All data and scripts are publicly available on GitHub for assay development (https://github.com/RobertsLab/resazurin-assay-development/releases/tag/v3.1) and the case study (https://github.com/RobertsLab/vims-resazurin/releases/tag/v1.0).

