## Supplementary Information for "From Blue to Pink: Resazurin as a High-Throughput Proxy for Metabolic Rate in Oysters"

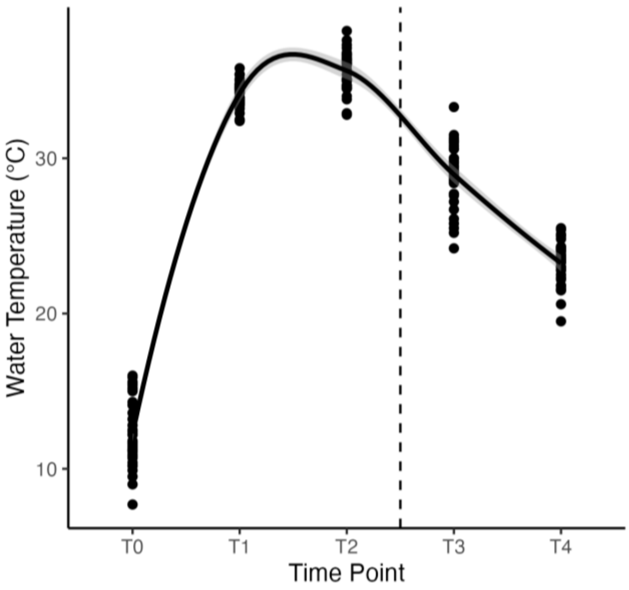


**Fig S1.** Temperatures (°C) recorded in plate wells during case study resazurin trials across hourly time points. The dashed line indicates the time at which plates were moved out of the incubator (40°C) for the recovery period.

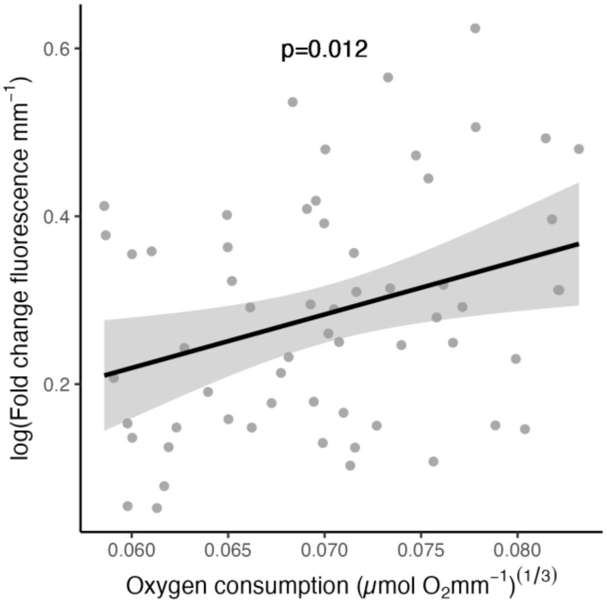


**Fig S2.** Correlation between oxygen consumption (μmol $O_2$ mm$^{-1}$; cube-root transformed) and fluorescence (fold change mm$^{-1}$; log-transformed) (y-axis) in oyster seed during 1 h measurements. Trendline indicates linear relationship between the two variables. P-value indicates significance of Pearson correlation test conducted on transformed fluorescence and oxygen consumption values.

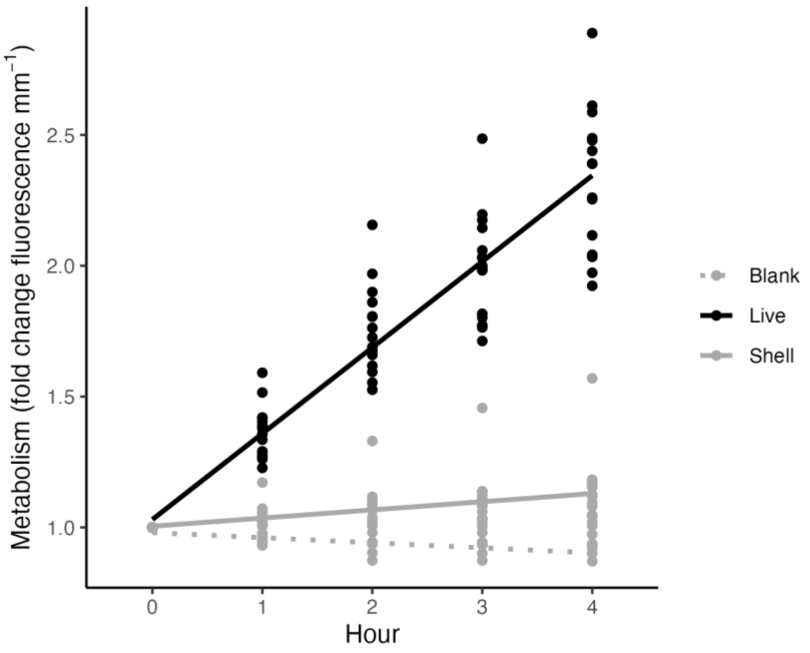


**Fig S3.** Metabolism (fold change in fluorescence) prior to size normalization in blank (dashed gray), empty shell (solid gray), and live oyster (solid black) samples.

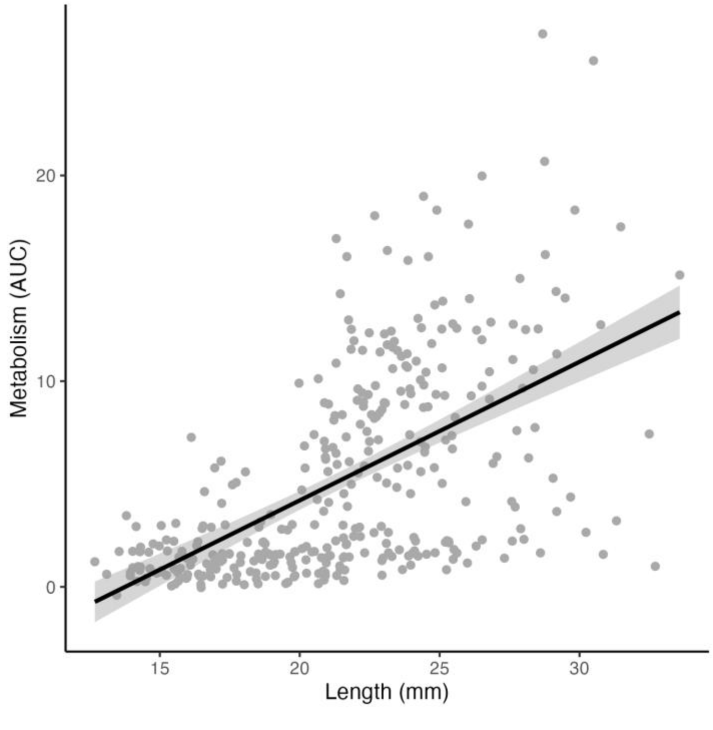


**Fig S4.** Correlation between shell length (mm) and metabolism (area under the curve; AUC) in oysters ranging from approx. 15-35 mm. Linear regression displays the relationship between size and AUC.

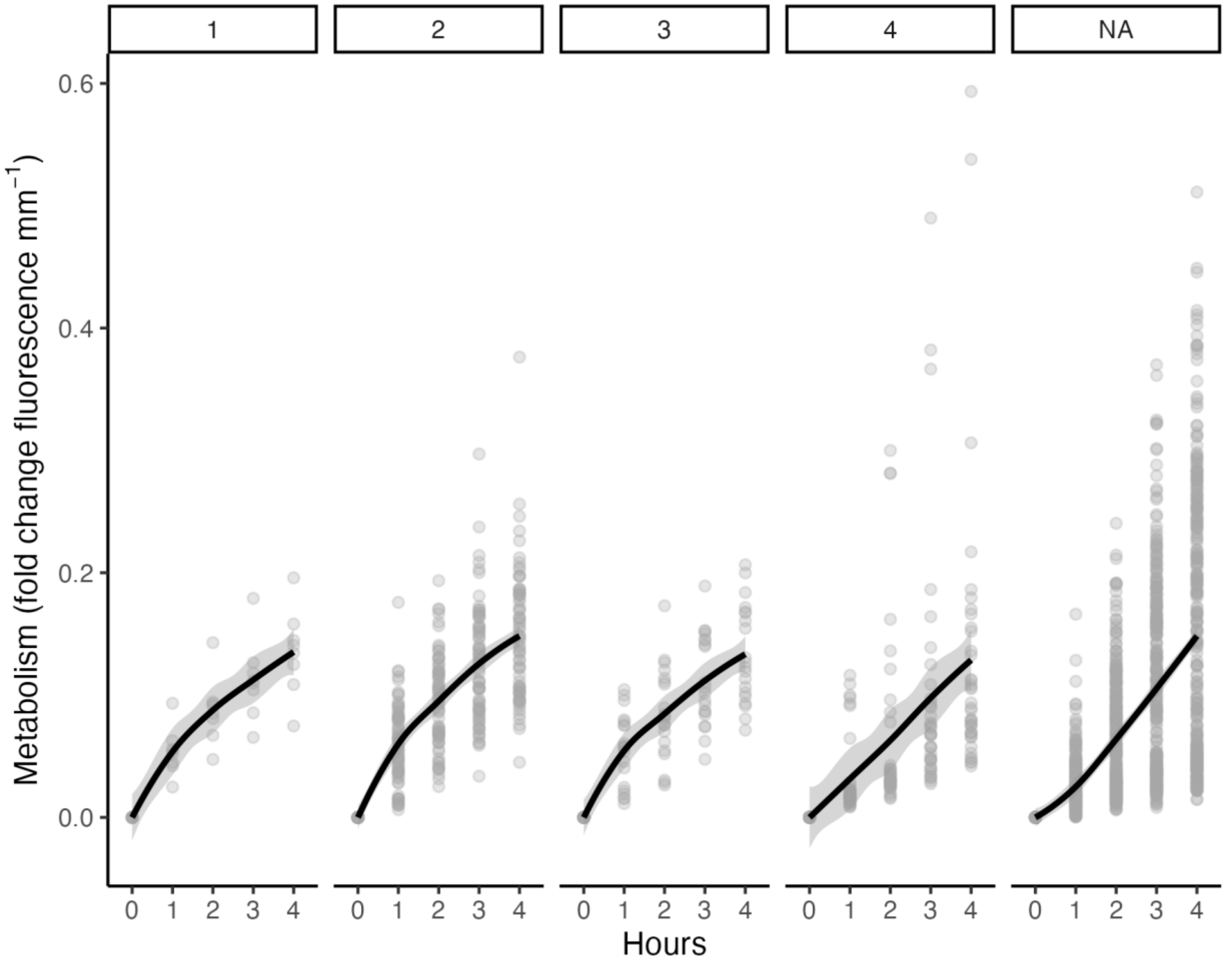


**Fig S5.** Metabolism (fold change in fluorescence mm$^{-1}$) for oyster seed (15-35 mm) depending on timing of mortality during a 4 h resazurin trial. Facet panels display time of mortality (1, 2, 3, or 4 h during a 4 h trial) with NA indicating oysters that remained alive for the entire trial. Lines show loess trend line with shading indicating standard error.

**Table S1.** Site selection for broodstock at low and high/moderate salinity sites.

| Site | Zone | Salinity (ppt) | Longitude | Latitude |
|---|---|---|---|---|
| Coan River, VA | Low salinity | 8 - 15 | 37.989 | -76.476 |
| Choptank River, MD | Low salinity | 6 - 13 | 38.593 | -76.129 |
| York River, VA | High/ moderate salinity | 18 - 23 | 37.248 | -76.501 |

**Table S2.** Temperature sensitivity (Q10) values of metabolism measured using resazurin fluorescence in oysters. Experiments were conducted in two size classes (3-8 mm and 5-15 mm). AUC indicates total metabolic activity calculated as area under the curve.

| Size class: 3-8 mm | | |
|---|---|---|
| **Temperature Interval** | **AUC** | **Q10** |
| 20-36°C | 4.11 | 1.22 |
| 36-42°C | 5.62 | 0.28 |
| **Size class: 5-15 mm** | | |
| **Temperature Interval** | **AUC** | **Q10** |
| 21-26°C | 2.14 | 1.14 |
| 26–32°C | 2.28 | 3.52 |
| 32–37°C | 4.86 | 0.55 |
| 37–42°C | 3.60 | 1.22 |
| 42–45°C | 3.98 | 0.01 |

### Appendix A: General Resazurin Protocol

This protocol describes the general procedure for conducting resazurin metabolic assays in oysters. The protocol will require modifications to fit the specific experimental design, hypotheses, and system.

See the GitHub page for assay development in the Roberts Lab at the University of Washington for more information and example data sets. https://github.com/RobertsLab/resazurin-assay-development

Resazurin metabolic assays are used to measure the metabolic rate of marine organisms. This protocol has been developed for use in oysters, but can be adapted for other organisms.

Standard Operating Protocol (SOP)

A. Overview

This protocol details general approaches for resazurin metabolic rate measurements in oysters. The protocol can be adapted for various sizes, temperature profiles, or different stress exposures following this base protocol.

B. Materials and preparing solutions

a. Containers

Here are our suggested vessels for conducting trials. The animals should be fully submerged and the container size will need to be scaled to the animal size appropriately.

- Small seed (<7mm length): trials conducted in 48 or 96 well plates
- Medium seed (15-40 mm length): trials conducted in 12 or 24 well plates or small plastic cups
- Large seed/adults (>40 mm length): trials can be conducted in tri-pour cups, beakers, or plastic cups. Scale volume appropriately.

We recommend performing preliminary trials to ensure that a change in resazurin fluorescence can be detected over the time scale desired.

b. Sample size

Plan your sample sizes appropriately depending on your research question. Metabolic rates are variable, so we recommend at least n=20 per experimental group as a minimum (higher replication is best).

When selecting containers and volume of solutions, remember to account for blanks (more information below). We recommend if you use plates that you include n=3-8 blanks per plate and if using cups, we recommend n=5-10 blanks per experimental treatment.

c. Stock resazurin solution

To make the saturated resazurin stock solution (10 mL) mix the following. This solution will be used for multiple trials. Scale the volume if required.

- 0.5 g resazurin salt (e.g., https://www.thermofisher.com/order/catalog/product/R12204)
- 10 mL deionized (DI) water
- 10 µL dimethyl sulfoxide (DMSO)

Store in a dark fridge or freezer.

d. Working resazurin solution

First, determine the volume of resazurin required depending on the size of your organism, the number of blanks and samples, and the container size as described above.

To prepare the working solution of resazurin, prepare the following.

- Desired working volume x 0.98666 = ___ mL filtered seawater (DI water with aquarium salts adjusted to appropriate salinity or filtered seawater)

- Desired working volume x 0.00222 = ___ mL resazurin stock solution as made above in step 1

- Desired working volume x 0.001 = ___ mL DMSO

- Desired working volume x 0.01 = ___ mL  antibiotic solution. We use 100x Penn/Strep & 100x Fungizone (https://us.vwr.com/store/product/4648458/null). This should be kept frozen in a dark freezer and thawed before use (thaw in the dark or cover with aluminum foil)

Store at 4°C in dark fridge until use. We recommend making a fresh batch of working stock and using within 5-7 days.

For example, here is a recipe for 150 mL of working stock.

- 148 mL seawater
- 333 µL resazurin stock solution
- 150 µL DMSO
- 1.5 mL antibiotic solution

C. Supplies

- Bench top incubators, shaking tables, or other material required for your desired stress or incubation treatments along with data loggers to record conditions
- Paper towels and bench paper/pads
- Tweezers, transfer pipettes, and forceps
- Dissecting microscope
- Spectrophotometer plate reader that detects in fluorescence mode and associated software
- Plate reader filters (if required) with excitation wavelength of 530 and emission wavelength of 590. For example, we are currently using an excitation 528 wavelength filter with a bandpass of 20 and an emission 590 wavelength filter with a bandpass of 20.
- Pipettes and tips
- Scale bar/ruler
- Camera/phone camera
- Plastic cups, beakers, plates, or other vessel for incubations

Label plates with identifying number (e.g. "Plate 1", "Plate 2") or label cups with unique numbers/identifiers.

D. Protocol

Conduct measurements at treatments desired for your experiments. For example, we typically conduct measurements at a control temperature and a high temperature. If multiple treatments are desired over multiple days, be sure to run a control treatment each day as reference and/or randomize the order of treatment conditions across days. Ensure you account for tank effects or other batch effects by randomizing loading order, position in incubators, etc.

For oysters, we have used the following treatments to detect metabolic responses to stress:

- Acute stress and survival testing: control temperature (10-20°C) and acute high temperature (36-42°C)
- Thermal performance testing: control temperature (10-20°C), and a gradient of temperatures from 25°C-45°C.
- Acute stress (40°C) for 2 hours followed by cooling/stabilizing temperatures (counter top or fridge for 2 hours)
- Measurements at non stressful temperatures to characterize metabolic rate variability between groups (e.g., 28°C)
- Measurements in mechanically stressed groups following shaking on shaker table.

We strongly recommend preliminary testing to determine appropriate temperature treatments for your study system and research question.

a. Time points

Resazurin measurements should be collected on a timescale appropriate for your research question. Typically, we run assays over a 3-6 hour period with measurements collected every 30-60 minutes. For example, here is a typical schedule for a day in which we are conducting 5 hour incubations at a control and high temperature.

08:00-09:00: Load plates with oysters, take size images, and load resazurin solution
09:00: Time 0 measurement
10:00: Time 1 measurement
11:00: Time 2 measurement
12:00: Time 3 measurement
13:00: Time 4 measurement
14:00: Time 5 measurement
14:00-16:00: Clean up and assess survival (if needed)

b. Load and prepare samples

Before starting, set the incubator at the desired temperature or set up your treatments.

1. Prepare animals for assays. Track the source tank, treatment, or other identifying information.
2. Add animals into labeled plates or containers and placing them into the empty container.
3. Add blanks into your design. These will be wells or cups loaded with only resazurin solution. See recommendations on number of blanks above.
4. Write the location of wells on a plate map if using plates.
5. Allocate the animals either into their designated cup, onto the lid of the plate, or on the counter.
6. Take images of each animal with a scale bar with their identifying information in the photograph.
7. Move the animals into their respective wells in the plate if you placed them on the lid or counter for a photograph.
8. Fill each well with the desired amount of resazurin working stock at ambient temperature using a microchannel pipette or graduated cylinder. The volume should be enough to fully submerge the animal without overflowing the container.

Measure size using ImageJ or other imaging software. We typically use maximum shell length (mm) for size normalization (see data analysis below).

c. Measurements

1. Turn on the computer and plate reader. Open the plate reader software.
2. Create a new protocol that conducts end point measurements from the top of the plate using an excitation wavelength of 530 and emission wavelength of 590 nm.

3. Name the protocol and save. This process may vary depending on your instrument.
4. Take a T0 initial measurement - this is critical! If using a plate, you can place the plate with the animals directly on the plate reader (do not have the lid on the plate). If you have animals in cups, take a small sample (250 µL) of the resazurin liquid from each container, place into a 96-well plate, and then conduct the measurements. If you use this method, be sure to make a plate map of the location of the samples and identifying information. Conduct measurements for blanks and samples.
5. Collect and export readings as directed in the plate software.
6. Save the file with a descriptive name. For example, `YYYYMMDD_TemperatureTreatment_Plate#_T0.xlsx`.
7. Save the data to a data repository or computer for analysis. We highly recommend platforming all data and analysis in R and GitHub.
8. Record the time of the measurement.
9. Repeat for any remaining plates or treatments.
10. Take temperature measurements in wells or cups if relevant.
11. Move the animals to their respective treatment conditions.
12. Repeat at 1, 2, 3, 4, and 5 hours of incubation or as necessary for your experiment.

d. Survival measurements

Either each hour (if you can easily see the animals in larger cups) or at the end of the incubation (for animals in 96 well plates), conduct survival assessments. Note that it is critical to perform survival assessments so that you can analyze resazurin metabolic response for those that survive and those that die during the trials. If performing trials in 96 well plates or other small containers, we recommend performing survival assessment at the end of the incubation. If you are conducting trials in larger cups where you can see the animals without removing the resazurin solution, you can assess survival at each time point.  This is likely only necessary if you are using acute stress treatments.

If using plates, prepare a plate map for recording the assessments - this is an easy method to keep track of the assessments. If you are not using plates, make a list of all samples with columns for recording survival at each time point.

If working with shellfish, use tweezers/forceps to take each animal out and examine in a petri dish filled with DI water under a dissecting scope. Determine if the oyster is dead by placing the cup side of the oyster down and gently taping/moving the shell. If the shell is open and remains open after tapping, the oyster is dead. If the shell is closed tight or closes after tapping, the oyster is alive. Use other determination methods for other organisms.

Generate a data frame that has columns for sample ID, treatments, date, and other relevant information. Add a column designated "mortality" and add a 0 for alive and 1 for dead animals. Edit as required for your specific analysis. See examples in our GitHub repository.

e. Waste disposal

Check with your Environmental Safety department to check on proper disposal procedures. At the University of Washington, for example, waste resazurin solution can be washed down the sink and flushed with plenty of water. Rinse containers thoroughly.

E. Data preparation and analysis

Prepare the following data frames (see examples at our repository):

- Size measurements: columns for sample ID, date, treatment, and size measurement (e.g., length in mm)
- Metadata: columns for sample ID, date, treatment, tank or batch effects, species, well/cup ID, and sample type (i.e., "blank" or "sample")
- Resazurin files: files exported from plate reader software that contain fluorescence readings for each well of the plate
- Mortality assessment (if applicable): columns for sample ID, date, treatment, and mortality assessment (e.g., 0 for alive and 1 for dead)

Conduct the following analysis steps (see R scripts available for use and adaptation in our repository):

- Read in data and combine into one data frame
- Normalize all fluorescence values to the initial time point (fluorescence at time X divided by fluorescence at time 0). Conduct normalization for both samples and blanks.
- Calculate the mean value for normalized blanks within each unit (e.g., mean of all blanks in each plate at each time point)
- Subtract the mean blank value from the fluorescence value of each sample from the respective unit and time point
- Size normalize the data by dividing fluorescence values by size of each sample
- This generates metabolism as fold change in fluorescence normalized to size
- Proceed with visualization and statistical analyses, including testing for effects of treatment or other effects of interest and examining metabolic differences between animals that lived and animals that died during the trials. See examples in our GitHub repository.
